# Horizontal gene transfer becomes disadvantageous in rapidly fluctuating environments

**DOI:** 10.1101/2020.08.07.241406

**Authors:** Akshit Goyal, David Gelbwaser-Klimovsky, Jeff Gore

**Author notes:** These authors contributed equally (Akshit Goyal, David Gelbwaser-Klimovsky).

## Abstract

Horizontal gene transfer (HGT) allows organisms to share genetic material with non-offspring, and is typically considered beneficial for evolving populations. Recent unexplained observations suggest that HGT rates in nature are linked with environmental dynamics, being high in static environments but surprisingly low in fluctuating environments. Here, using a geometric model of adaptation, we show that this trend might arise from evolutionary constraints. During adaptation in our model, a population of phenotype vectors aligns with a potentially fluctuating environmental vector while experiencing mutation, selection, drift and HGT. Simulations and theory reveal that HGT shapes a trade-off between the adaptation speed of populations and their fitness. This trade-off gives rise to an optimal HGT rate which decreases sharply with the rate of environmental fluctuations. Our results are consistent with data from natural populations, and strikingly suggest that HGT may sometimes carry a significant disadvantage for populations.

---

Horizontal gene transfer (HGT), by which organisms exchange genetic material with non-offspring, is a fundamental force in microbial evolution [1–4]. The ability to accept and incorporate the DNA of others can often help asexual organisms such as bacteria acquire new traits from their neighbors, and is thus generally assumed to be beneficial for adapting populations [5, 6]. For example, a new metabolic pathway need not independently evolve in multiple individuals; it can instead evolve in one individual and spread to others via HGT [7–9].

The apparent advantages of HGT for adaptation suggest that it should be common in bacterial populations. Indeed, bioinformatic analyses suggest that HGT rates in bacteria can often be as high as 100 times their mutation rates, consistent with recent theory suggesting that higher HGT rates are generally more adaptive [10–14]. Surprisingly however, recent estimates show that HGT rates can vary widely based on the nature of the environment — being typically high in static environments like the ocean (~100x compared with mutation), but remarkably low in rapidly fluctuating environments like soils (~1x compared with mutation) [15–18]. This surprising but systematic variability in HGT rate remains poorly understood, but suggests that the advantage of horizontal gene transfer is linked with environmental dynamics.

Here, through a minimal model of an evolving population, we show that horizontal gene transfer can become increasingly maladaptive in rapidly fluctuating environments. In our model, individuals are phenotype vectors which rotate due to mutation and HGT, and eventually align with an environmental vector representing the optimal (fittest) phenotype. By studying the evolutionary dynamics of populations with different HGT rates in both static and fluctuating environments, we gain the following insight about why HGT can be maladaptive — in our model, HGT shapes a trade-off between the adaptation speed and fitness (once adapted) of populations. Namely, populations with higher rates of HGT are ultimately fitter than those with lower rates, but take longer to get to their highest fitness. As a consequence, for every environmental fluctuation rate, there is an optimal (or fittest) HGT rate which decreases with the rate of environmental fluctuations. In relatively static environments with low fluctuation rates, adaptation speed is less important than the maximum fitness because environmental changes are slow; this makes high HGT rates fitter. In contrast, in rapidly fluctuating environments, populations do not reach their maximum fitness and adaptation speed becomes the key factor determining evolutionary success, rendering populations with low HGT rates fitter.

Taken together, our results suggest how horizontal gene transfer may sometimes carry a significant disadvantage for evolving populations, and mechanistically explain how HGT rates in nature can be linked with environmental dynamics.

## RESULTS

### A vector-based model of an evolving population with mutation and horizontal transfer

Our model follows a population of asexually reproducing individuals evolving in response to a potentially fluctuating environment. Each individual is a vector, whose fitness depends on its alignment with another vector representing the environment (Fig. 1a). Evolutionary processes can influence these vectors in different ways. Mutation of an individual results in a small rotation of its vector in a random direction (Fig. 1b, top). Horizontal gene transfer between a donor and acceptor results in the acceptor’s vector rotating towards the donor’s (Fig. 1b, middle). And every generation, individual vectors produce offspring, making copies of themselves (Fig. 1b, bottom). Due to selection by the environment, fitter individuals have more off-spring. We will describe the ingredients of the model in this section, and detail the evolutionary processes in the next section.

**FIG. 1.**
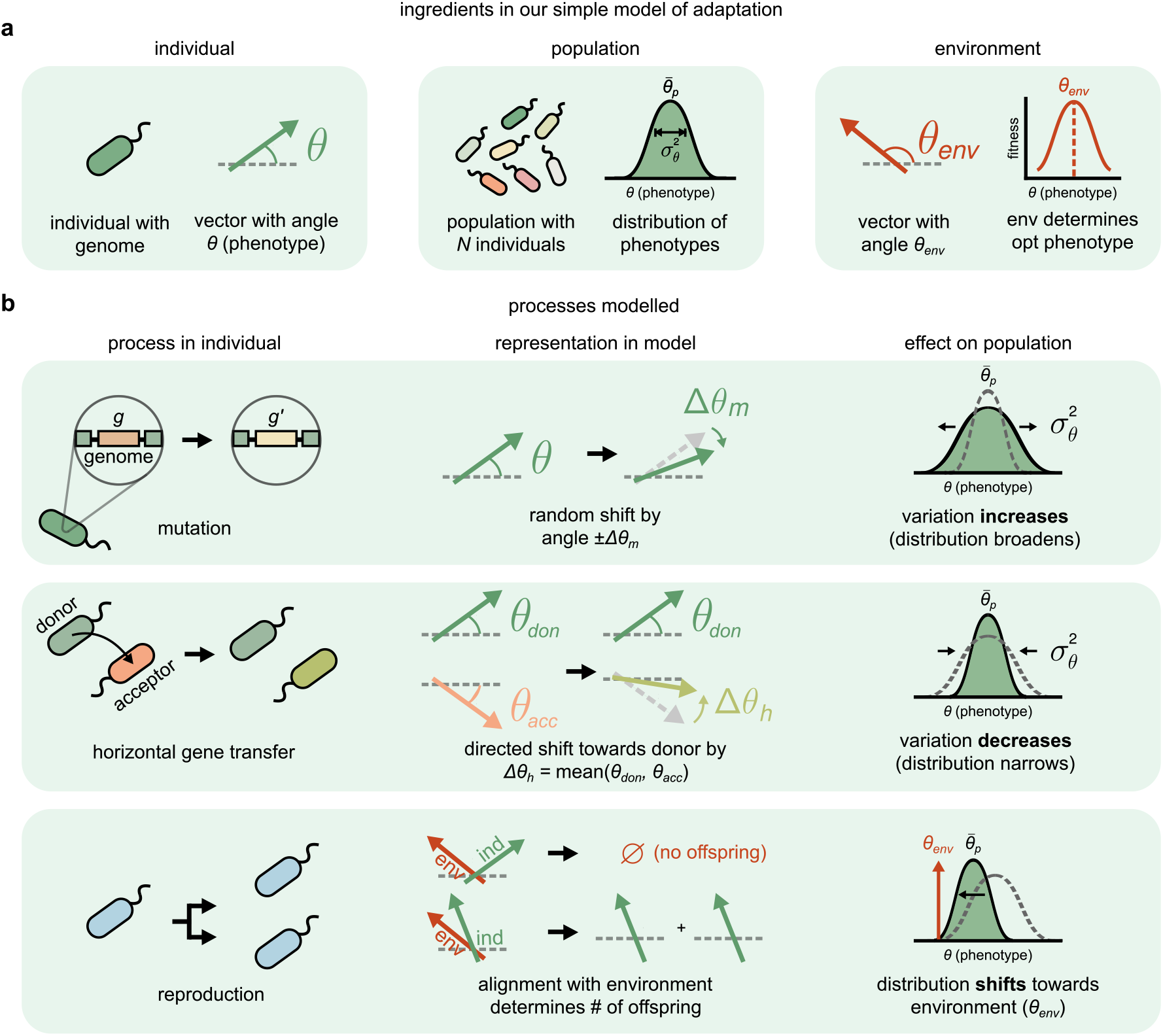
A vector-based model of an evolving population. **a,** [left] In our model, an asexually reproducing individual microbe is represented as a vector (green arrow) with unit length and at an angle *θ* representing its overall (or most important) phenotype. Evolutionary processes rotate this angle *θ* along a circle (0 ≤ *θ* < 2*π*). [middle] A population of *N* individuals is represented by a population of *N* vectors, characterized by a normal distribution of phenotypes, with mean 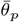 and variance 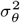. [right] The environment (red arrow) is represented as another vector at angle *θ*_env_, representing the optimal (or fittest) phenotype (red curve shows fitness landscape). **b,** Evolutionary processes in the model: [top] Mutation in an individual changes a gene *g* to *g*′. In the model, mutation rotates an individual vector in a random direction by Δ*θ*_*m*_. After several mutations in a population, 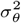 increases without changing 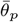. [middle] Horizontal gene transfer (HGT) between a donor and acceptor individual brings the acceptor’s phenotype closer to the donor’s. HGT rotates the acceptor’s vector towards the donor by Δ*θ*_*h*_. After several HGTs, 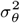 decreases without changing 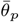. [bottom] In the model, individual vectors (green) produce offspring proportional to their fitness, i.e., their alignment with the environment (red). Reproduction shifts 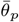 towards *θ*_env_ and reduces 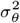.

In nature, the phenotype of an individual (its set of traits) depends on its genome in a complex manner [19]. For instance, an individual’s overall phenotype may consist of its ability to metabolize the available nutrients, defend against toxins, and maintain osmotic balance. For simplicity, our model coarse-grains these details: each individual is instead represented by a vector with angle *θ*, which represents its overall (or most important) phenotype (Fig. 1a, left). The phenotype vector of an individual can take any continuous value between 0 and 2*π*, but always has length 1. We assume each vector rotates in a circle, such that *θ* = 0 and *θ* = 2*π* represent the same angle or phenotype. Since each individual is a phenotype, a population of *N* individuals can be thought of as a distribution of phenotypes, with mean 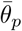 and variance 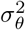 (Fig. 1a, middle). While we imagine each individual to be a bacterium, our model is general and applies to asexual organisms (such as other microbes) which experience mutation and horizontal gene transfer.

We also model the environment as a vector, with angle *θ*_env_, representing the optimal (or fittest) phenotype (Fig. 1a, right). The fitness of an individual is thus the alignment between its vector and the environment. Perfectly aligned individuals have the highest fitness, while those with vectors farther from the environment have lower fitness (Fig. 1a, right). Mathematically, an individual with phenotype *θ* has fitness 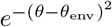. Using other fitness functions, as long as they smoothly decrease away from the optimum, does not affect our main conclusions (Fig. S1). Note that the environment *θ*_env_ may change over time, and does need not to be constant. This is how we will study fluctuating environments later in the text.

Our model shares some preliminary similarities with Fisher’s geometric model (FGM), specifically our representation of individuals as phenotypes and the environment as an optimal phenotype [20–23]. However, in addition to several technical differences between our vector-based model and FGM (Appendix 1), we emphasize that FGM studies the fitness effects of mutations alone; it lacks any notion of HGT and environmental fluctuations, which form the basis of our manuscript. Unlike mutations, which occur independently in individuals, HGT generates correlations between them.

### Evolutionary dynamics in the model

Evolutionary dynamics in our model proceed in discrete steps. At each step, one of three processes occurs. First, individuals mutate, each with a probability *p*_*m*_. In our model, a mutation causes an individual’s phenotype to shift randomly in either direction by Δ*θ*_*m*_ (Fig. 1b, top). Over time, the accumulation of mutations broadens the population’s phenotype distribution, increasing its variance without changing its mean (Fig. 1b, top right).

Second, pairs of individuals engage in horizontal gene transfer. In our model, each individual has a probability *p*_*h*_ of being part of an HGT pair. Each pair consists of a donor and an acceptor, assigned randomly. A horizontal transfer causes the acceptor’s phenotype (*θ*_acc_) to shift towards the donor’s phenotype (*θ*_don_) by an amount Δ*θ*_*h*_, which depends on *θ*_don_ and *θ*_acc_ (Fig. 1b, middle). For simplicity, Δ*θ*_*h*_ = mean(*θ*_don_*, θ*_acc_), but as long as the acceptor’s phenotype moves in the direction of the donor, the magnitude of Δ*θ*_*h*_ does not affect our main conclusions (Appendix 1). This choice is inspired by the effect of the transfer of a genetic segment from a donor to an acceptor organism: namely that after the transfer, the acceptor tends to acquire a phenotype more similar to the donor [12, 24–26]. HGT does not always benefit individuals; it depends on which genes are gained, and in which environment [27]. Our model captures this: an HGT event can both increase and decrease the fitness of an acceptor, depending on whether the acceptor is more aligned with the environment than the donor. Over time, several HGT steps narrow the phenotype distribution, reducing its variance without changing its mean (Fig. 1b, middle right). Note that this is a statement about phenotypic variance, not genetic variance.

Finally, individuals in the population reproduce. We model reproduction similar to a Wright-Fisher process, and account for both fitness differences between individuals (selection) and stochasticity in birth-death events (drift). During reproduction, we determine the next generation of individuals by sampling individuals from the current generation with a probability proportional to their fitness (Fig. 1b, bottom). For simplicity, we keep the population size fixed. Reproduction shifts the mean of the phenotype distribution towards the environment, *θ*_env_ (Fig. 1b, bottom right).

Using these three processes, we simulate evolutionary dynamics generation by generation. At the beginning of each simulation, we generate a population of *N* = 5, 000 individuals whose phenotypes are normally distributed. Increasing the population size does not affect our results (Appendix 1). At every time step, the population either mutates or undergoes horizontal gene transfer. After every generation (1, 000 time steps) the population reproduces. In the rest of the manuscript, we measure time in units of generations. In each simulation, we control the HGT rate of a population, relative to mutation, by changing the probabilities *p*_*m*_ and *p*_*h*_, the respective intensities of mutation and HGT per step. We henceforth refer to *p*_*m*_ and *p*_*h*_ as the mutation and HGT rates, respectively. Note that the specific parameter values we used here do not impact our conclusions qualitatively (values listed in Appendix 5).

### HGT shapes a trade-off between fitness and adaptation speed

In order to understand the effects of HGT rates on adaptation, we first analyzed adaptation in static environments. For this, we simulated the evolutionary dynamics of two populations: one with a lower HGT rate; the other with a higher HGT rate. At the beginning of each simulation (time *t* = 0), both populations were otherwise identical — they both had phenotype distributions with mean 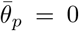 and variance 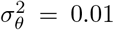. The environment of both populations was arbitrarily positioned at a *θ*_env_ = 1 (dashed line in Fig. 2a). The population with low HGT rate had *p*_*h*_ = 0.01, while the one with high HGT rate had *p*_*h*_ = 0.1; both populations had identical mutation rates *p*_*m*_ = 0.01. In what follows, we will refer to the process of a population aligning with the environment as adaptation; adapted populations will be those whose mean phenotypes have successfully aligned 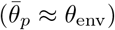.

**FIG. 2.**
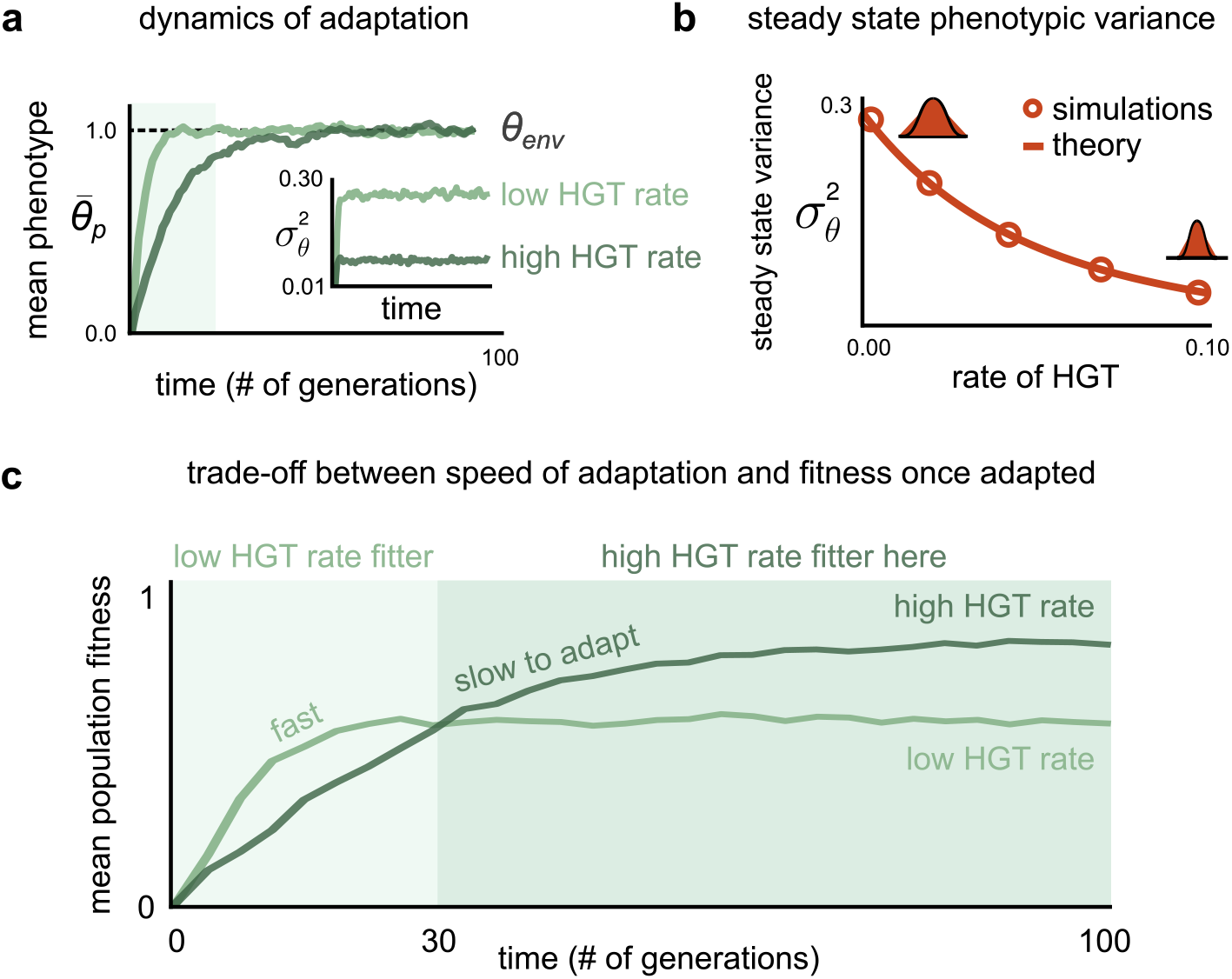
HGT shapes a trade-off between fitness and adaptation speed. **a,** Time evolution of the mean phenotype 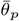 of two evolving populations: one with high HGT rate (dark green; *p*_*h*_ = 0.1), the other with low HGT rate (light green; *p*_*h*_ = 0.01). Both populations were otherwise identical, with mutation rate *p*_*m*_ = 0.01, and initial conditions (at *t* = 0, 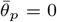 and variance 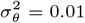). The dashed black line shows the environment *θ*_env_ = *π*, which was static. The inset shows the time evolution of the phenotype variance 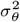. **b,** The phenotypic variance at steady state as a function of the rate of HGT of a population. The open circles show results from simulations of our model, while the line shows results from an analytical derivation (Appendix 1). **c,** Time evolution of the average fitness for both the populations in **a**. Background colors represent two adaptive regimes: at early (0 < *t* ≤ 30) and late times (*t* > 30) (see Results).

As expected, we found that after a sufficiently long time (100 generations) both populations had reached a steady state and were adapted, i.e., the means of their phenotype distributions, 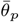 had aligned with the environment, *θ*_env_ (Fig. 2a). However, the variances of both phenotype distributions, which had also reached steady state, were quite different. Specifically, the variance of the population with the higher HGT rate was roughly 3-fold lower (Fig. 2a, inset). This suggested a general negative relationship between the steady state phenotypic variance 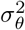 and the HGT rate of a population in our model, which we confirmed by repeating these simulations for populations at several other HGT rates (Fig. 2b). We also analytically derived this relationship, and found that it closely agreed with simulations (Fig. 2b, open circles).

Another difference between the adaptive dynamics of both populations was their speeds — populations with high HGT rates were slower and took longer to get to steady state (Fig. 2c). This is because when not adapted, the lower the variance, the lower is the chance that any individual is aligned with or near the environment, resulting in a smaller shift per generation. Additionally, since high HGT rate populations had lower phenotypic variance, at steady state their individuals were fitter on average. This is because once adapted, the lower the variance, the closer an individual is to the optimal phenotype, *θ*_env_ (Fig. 2c). Therefore, increasing HGT rate affects a population in two different ways: it decreases its adaptation speed but increases it fitness at steady state. In this way, HGT shapes a trade-off between adaptation speed and fitness.

To understand the consequences of such a trade-off, consider the two example adaptive trajectories illustrated in Fig. 2c. These trajectories can be partitioned into two time regimes: an early regime (0 < *t* ≤ 30) and a late regime (*t* > 30). In the early regime, the low HGT rate population adapts faster and is hence fitter, while over longer times, the high HGT rate population gets enough time to adapt and eventually becomes fitter instead. This qualitative difference suggests that the answer to which HGT rate is fitter would depend on the typical time a population has to adapt to its environment, say *τ*_env_. This observation prompted us to explore adaptive dynamics in fluctuating environments, which we describe in the next section.

### The optimal HGT rate decreases sharply with the rate of environmental fluctuations

To better understand the link between HGT rates and environmental dynamics in our model, we extended it to include fluctuating environments. In contrast with static environments, where the optimal phenotype *θ*_env_ stayed fixed (Fig. 3a, top), in fluctuating environments, after a characteristic time period *τ*_env_, *θ*_env_ abruptly changed to a random optimal phenotype 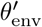 (Fig. 3a, bottom). For simplicity, we picked 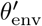 randomly, though we verified that introducing a characteristic magnitude of environmental changes Δ*θ*_env_ did not affect our results (Fig. S3).

**FIG. 3.**
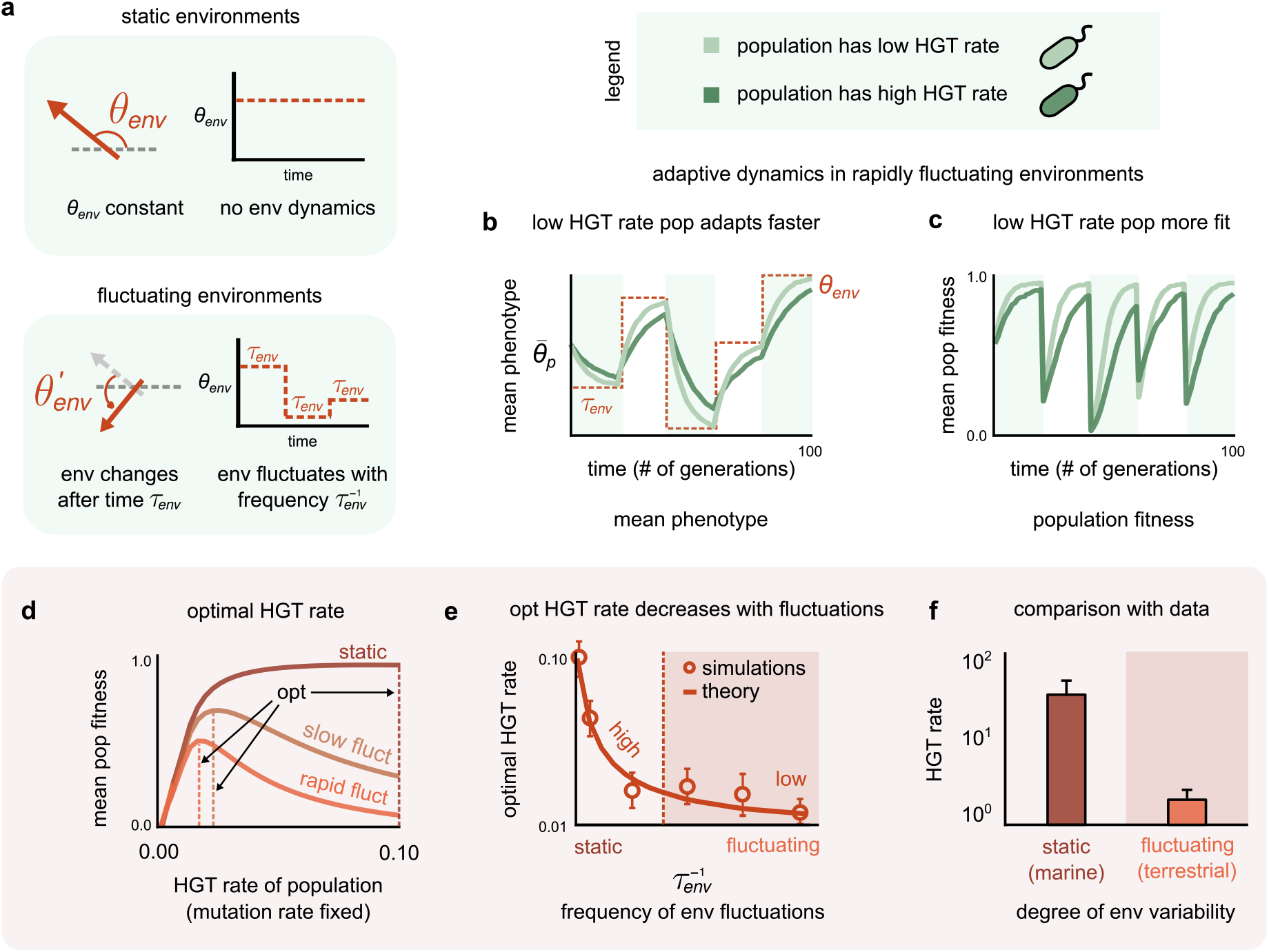
The optimal HGT rate decreases sharply with the rate of environmental fluctuations. **a,** In our model, a static environment is represented as a fixed vector at angle *θ*_env_, which does not change with time. A fluctuating environment changes to a random angle 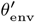 after time *τ*_env_ (which we modulate). **b,** Time evolution of the mean phenotype 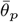 and **c,** mean fitness of two evolving populations in a rapidly fluctuating environment (dashed line; 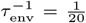). Both populations are initially identical, except one has high HGT rate (dark green; *p*_*h*_ = 0.1), while the other has low HGT rate (light green; *p*_*h*_ = 0.01). **d,** Curves showing the average fitness of an individual in a population as a function of its HGT rate *p*_*h*_ (with fixed mutation rate *p*_*m*_ = 0.01). Each curve represents a unique environmental fluctuation frequency 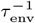: static (brown;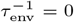), slowly fluctuating (mustard; 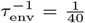) and rapidly fluctuating (orange; 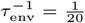). Dashed lines represent the optimal HGT rate for that 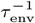. **e,** The optimal HGT rate as a function of 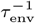. The open circles show results from simulations of our model, while the line shows results from an analytical derivation (Appendix 1). Error bars represent standard deviation. **f,** Bar plots of the average HGT (specifically, homologous recombination) rates measured relative to mutation using microbial genomes from static, marine environments (brown) and fluctuating, terrestrial environments (orange) (data from ref. [15]). Values represent mean; error bars represent standard deviation.

Following our analysis in static environments, we simulated the adaptive dynamics of two different populations in a fluctuating environment. As before, both populations differed only in their HGT rates: one was high, the other low (*p*_*h*_ = 0.1 and 0.01 respectively). In rapidly fluctuating environments (at high fluctuation frequency 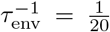 per generation), we found that the population with a low HGT rate both adapted to the environment quicker (Fig. 3c), and was typically fitter than that with a high HGT rate (Fig. 3d). This is best explained through the two adaptive regimes in Fig. 2c. In static or slowly fluctuating environments, the time between fluctuations is long; thus populations are mostly in the late regime, in which the population with the higher HGT rate is fitter. However, as environments fluctuate more rapidly, populations become restricted to the early regime, in which the high HGT rate population cannot reach its highest fitness; this makes the quicker adapting, or low HGT rate population fitter.

Thus, the fitness of individuals in a population depends on both the HGT rate and the degree of environmental fluctuations. To systematically probe this dependence, we extended our simulations to incorporate several populations with different HGT rates and environmental fluctuation frequencies (HGT rates ranged from *p*_*h*_ = 0.0 to 0.1; fluctuation frequencies from 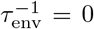 (static) to 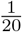 per generation (rapidly fluctuating)). For each environmental fluctuation frequency, we calculated the mean fitness of individuals in a population over time with the population’s HGT rate. We found that in each environment, a population with a specific HGT rate was fittest (Fig. 3d). We called each such HGT rate the optimal HGT rate for that frequency of environmental fluctuations. By extracting these optimal HGT rates for every environmental fluctuation frequency, we found that the optimal rate of HGT decreased sharply — by an order of magnitude in our model — with increasing environmental fluctuations (Fig. 3e). Theoretical calculations confirmed this result (Fig. 3e, open circles), and explained that the optimal HGT rate was one that optimized the phenotypic variance of the population for a specific environment. Having an optimal variance allows a population to achieve a balance between speed and ultimate fitness, crucial in fluctuating environments (Appendix 1).

### Comparison with data

The sharp decrease in optimal HGT rate observed in our model provides a possible explanation for a specific trend in HGT rates observed in natural microbial populations. Specifically, our model shows that while high HGT rates are optimal in static environments, they become increasingly suboptimal in fluctuating environments. Others have suggested, through a comparison of abiotic and biotic factors, that terrestrial environments are relatively fluctuating while marine environments are relatively static [16]. Consistent with our model, a recent study showed that microbial genomes from marine environments have on average high HGT rates (> 1; average 27.9 ± 3.2), while those from terrestrial environments have on average low HGT rates (~1; average 1.5 ± 0.6; Appendix 3) [15]. Here, the HGT rate of each microbial genome was measured as the homologous recombination rate relative to mutation. Further, endosymbiont genomes such as *Wolbachia*, whose life cycles occur inside relatively static environments such as host cells, also have high HGT rates (> 1; ~ 5.5 ± 0.5) [28].

## DISCUSSION

Here, we have attempted to explain the origin of a surprising relationship between horizontal gene transfer (HGT) rates and environmental dynamics that is observed in nature. Specifically, HGT rates are typically high in relatively static environments and low in relatively fluctuating environments [15, 16]. A corollary of this relationship is the following counterintuitive statement: high rates of horizontal gene transfer are disadvantageous for populations (maladaptive) in fluctuating environments. Our explanation is grounded in a vector-based minimal model of a population evolving via mutation, HGT, selection and drift, which we propose and analyze in this manuscript. Key to our explanation is a trade-off between adaptation speed and maximum fitness that our model elucidates. Specifically, a high rate of horizontal transfer brings the phenotypes of individuals in a population closer to each other, reducing its phenotypic variance. A decreased phenotypic variance helps a population localize better around fitness peaks, but also prolongs its search for those peaks. This trade-off manifests as an optimal HGT rate for every environmental fluctuation frequency. We find that the optimal HGT rate decreases sharply with environmental fluctuations, consistent with existing data on microbial recombination rates (Figs. 3e and 3f, respectively).

Our comparison of HGT rates from natural populations relied on the assumption that terrestrial environments are relatively fluctuating, while marine environments are relatively stable. While this is a simplification, others have argued that terrestrial environments are indeed typically more fluctuating; these arguments are based on both abiotic factors such as salinity, pH and oxygen availability, as well as biotic factors such as the size of microbial regulatory networks (which has been shown to correlate with environmental variability) [16]. The development of more rigorous criteria may be needed to more accurately classify environments as static or fluctuating.

Further, we understand that by confining our analysis to an isolated population, our model ignores the migration of individuals from other populations, which can often bring new beneficial genetic material from an external reservoir of genes [29]. More importantly, unlike HGT between individuals of the same population, HGT with individuals of an external population may increase phenotypic variance rather than decrease it. In this manuscript, we refrained from a full exploration of the effects of migration. However, we verified that, even in an extreme case, migration was unlikely to qualitatively affect our main conclusions. For this “worst-case” analysis, we added migration from an external population that was always perfectly adapted, i.e., its individuals were always artificially aligned with the internal population’s environment. We found that even in this extreme scenario, the optimal HGT rate decreased with environmental fluctuations (Fig. S2).

Throughout this study, we kept mutation rates fixed, thus implicitly assuming that HGT rates are more plastic than mutation rates. Data suggest that this is a reasonable assumption, and is often used while inferring HGT rates from microbial genomes [15, 30, 31]. Nevertheless, some of our observations apply to mutation rates as well. For instance, by increasing the phenotypic variance, mutation may also shape the trade-off between adaptation speed and fitness.

Finally, despite its simplicity, our vector-based model captures several evolutionary phenomena exhibited by traditional population genetics models, such as mutation-selection balance (Fig. S4) and Muller’s ratchet (Fig. S5). We believe this makes it a useful tool to study the interplay between mutation and HGT during evolution. Due to its simplicity, we can also extend the model to study the effects of HGT in other scenarios. For instance, we can study the impact of HGT on the co-evolution between hosts and pathogens (such as bacteria and phage) which constantly modify each other’s fitness landscapes; as well as design protocols that help reduce the fitness of certain bacterial species with high HGT rates, such as antibiotic resistant bacteria. These remain promising directions for future research.

## Acknowledgements

We thank Shaul Pollak, Rohini Subrahmanyam, Anjali Jaiman, Natalie Vaisman and members of the Gore lab for valuable discussions. A.G. and D.G.K. are supported by the Gordon and Betty Moore Foundation as Physics of Living Systems Fellows through grant number GBMF4513.

