## Supplemental Material for "Horizontal gene transfer becomes disadvantageous in rapidly fluctuating environments"

Akshit Goyal, David Gelbwaser-Klimovsky and Jeff Gore
*Physics of Living Systems, Department of Physics,*
*Massachusetts Institute of Technology, Cambridge, MA 02139, USA.*

#### APPENDIX 1
#### DERIVATION OF STEADY STATE PHENOTYPIC DISTRIBUTIONS

To understand the analytical nature of the relationship between the steady state phenotypic variance and the HGT rates in our model, we used a simplified version of it. The key simplification was that all evolutionary processes preserve the shape of the phenotype distribution in the population. The relatively good agreement between the simulations and the theoretical results showed that this is a good approximation. A population whose phenotypes are initially normally distributed thus remains a population with normally distributed phenotypes. All evolutionary processes therefore only change the mean $\bar{\theta}_p$ and variance $\sigma_\theta^2$ of the population's phenotype distribution. This theory allows us to approximate all the relevant quantities in our model. We now follow each of the evolutionary processes step by step, and describe the effect of each step on the population's phenotype distribution. Using these effects, we will later derive an expression for the steady state phenotypic variance, after several generations.

Throughout this section, consider that the original phenotype distribution $\mathcal{P}(\theta)$ is a Gaussian with mean $\bar{\theta}_p$ and variance $\sigma_\theta^2$, given by the following expression:

$$\mathcal{P}(\theta) = \frac{1}{\sqrt{2\pi}\sigma_\theta} e^{-\frac{(\theta-\bar{\theta}_p)^2}{2\sigma_\theta^2}} \tag{S1}$$

##### Horizontal gene transfer

In our model, during a horizontal gene transfer (HGT) step, two individuals with a probability $p_h$ engage in horizontal gene transfer, HGT). Consider two such individuals with phenotypes $\theta_1$ and $\theta_2$. One of these, at random, is the donor — whose phenotype does not change during HGT — and the other the acceptor, whose phenotype changes to the mean phenotype $\frac{\theta_1+\theta_2}{2}$. Therefore, during each HGT event, only half of the participants' phenotypes change. During an HGT step, a fraction $\frac{p_h}{2}$ of the phenotype distribution, corresponding to the total acceptor population changes; this change is given by the following expression:

$$\frac{p_h}{2}\int \frac{1}{2\pi\sigma_\theta^2} e^{-\frac{(\theta_1-\bar{\theta}_p)^2}{2\sigma_\theta^2}} e^{-\frac{(\theta_2-\bar{\theta}_p)^2}{2\sigma_\theta^2}} \delta\Big(\frac{\theta_1+\theta_2}{2}-\theta\Big) d\theta_1 d\theta_2 = \frac{p_h}{2}\frac{1}{\sqrt{\pi}\sigma_\theta} e^{-\frac{(\theta-\bar{\theta}_p)^2}{\sigma_\theta^2}} \tag{S2}$$

Here, $\delta$ represents the Kronecker delta, which is $\delta(x) \to \infty$ when $x = 0$ and $\delta(x) = 0$ everywhere else, i.e., $x \neq 0$. The remaining fraction, $1 - \frac{p_h}{2}$ of the distribution, does not change. The distribution $\mathcal{P}'(\theta)$ after an HGT step is therefore:

$$\mathcal{P}'(\theta) = (1-\frac{p_h}{2})\frac{1}{\sqrt{2\pi}\sigma_\theta} e^{-\frac{(\theta-\bar{\theta}_p)^2}{2\sigma_\theta^2}} + \frac{p_h}{2}\frac{1}{\sqrt{\pi}\sigma_\theta} e^{-\frac{(\theta-\bar{\theta}_p)^2}{\sigma_\theta^2}} \tag{S3}$$

Therefore, using equation (S3), we see that after an HGT step, the mean phenotype does not change, but the phenotypic variance changes. The variance $\sigma_\theta'^2$ after an HGT step is given by the following expression:

$$\sigma_\theta'^2 = (1-\frac{p_h}{2})\sigma_\theta^2 + \frac{p_h}{4}\sigma_\theta^2 \tag{S4}$$

$$= \sigma_\theta^2(1-\frac{p_h}{4}). \tag{S5}$$

We assume that after an HGT step, the phenotype distribution is still a Gaussian with mean $\theta_p$ and variance $\sigma_\theta'^2$.

**Mutation**

To calculate the population's phenotypic distribution after a mutation step, we follow a similar procedure to that for HGT. During a mutation step, a fraction $p_m$ of individuals in the population mutate, and as a result of each mutation, a mutant's phenotype changes in a random direction by an amount $\Delta\theta$. This implies that we assume that all mutational effects have the same magnitude of effect, which is done for simplicity (i.e., one can show that our results will not qualitatively change if one considers more realistic distributions of mutational effect). On average, half the mutants will have their phenotype values increase by $\Delta\theta$, and half will have their phenotype values decrease by $\Delta\theta$. Using these assumptions, we can calculate the phenotype distribution after a mutation step $\mathcal{P}'(\theta)$, and show that it is given by the following expression:

$$\mathcal{P}'(\theta) = (1-p_m)\frac{1}{\sqrt{2\pi}\sigma_\theta}e^{-\frac{(\theta-\bar{\theta}_p)^2}{2\sigma_\theta^2}} + \frac{p_m}{2}\Big(\frac{1}{\sqrt{2\pi}\sigma_\theta}e^{-\frac{(\theta+\Delta\theta-\bar{\theta}_p)^2}{2\sigma_\theta^2}} + \frac{1}{\sqrt{2\pi}\sigma_\theta}e^{-\frac{(\theta-\Delta\theta-\bar{\theta}_p)^2}{2\sigma_\theta^2}}\Big).$$

Similar to HGT, we see that a mutation step does not change the mean of the distribution, but increases its variance ${\sigma'_\theta}^2$, which after a mutation step, is related to the variance $\sigma_\theta^2$ before the step as follows:

$${\sigma'_\theta}^2 = \sigma_\theta^2 + p_m\Delta\theta^2. \tag{S6}$$

We assume that after an HGT step, the phenotype distribution is still a Gaussian with mean $\theta_p$ and variance ${\sigma'_\theta}^2$.

**Reproduction**

During reproduction, the population undergoes selection, using a fitness function given by each individual's alignment with the environment. Consider, for simplicity, that the fitness landscape is itself a Gaussian function with a peak $\theta_{\text{env}}$ and a variance $\sigma_{\text{env}}^2$. After a round of selection, the population's phenotype distribution $\mathcal{P}'(\theta)$ is proportional to the following:

$$\mathcal{P}'(\theta) \propto \frac{1}{\sqrt{2\pi}\sigma_\theta}e^{-\frac{(\theta-\bar{\theta}_p)^2}{2\sigma_\theta^2}}\frac{1}{\sqrt{2\pi}\sigma_{\text{env}}}e^{-\frac{(\theta-\theta_{\text{env}})^2}{2\sigma_{\text{env}}^2}} \tag{S7}$$

After rearranging terms, we get the following expression for $\mathcal{P}'(\theta)$:

$$\mathcal{P}'(\theta) \propto \exp\Big[\frac{-\big(\theta - \frac{\bar{\theta}_p\sigma_{\text{env}}^2+\theta_{\text{env}}\sigma_\theta^2}{\sigma_{\text{tot}}^2}\big)^2}{\frac{2\sigma_\theta^2\sigma_{\text{env}}^2}{\sigma_{\text{tot}}^2}}\Big], \tag{S8}$$

where $\sigma_{\text{tot}}^2 = \sigma_\theta^2 + \sigma_{\text{env}}^2$. After a reproduction step, we see that population's mean phenotype changes, and the new mean $\bar{\theta}'_p$ is given by:

$$\bar{\theta}'_p = \frac{\bar{\theta}_p\sigma_{\text{env}}^2 + \theta_{\text{env}}\sigma_\theta^2}{\sigma_{\text{tot}}^2} \tag{S9}$$

The difference between the new mean phenotype and the optimal phenotype $\theta_{\text{env}}$ is as follows:

$$\bar{\theta}'_p - \theta_{\text{env}} = \frac{\sigma_{\text{env}}^2}{\sigma_{\text{tot}}^2}(\bar{\theta}_p - \theta_{\text{env}}) \equiv \frac{1}{\Sigma_t^2}(\bar{\theta}_p - \theta_{\text{env}}). \tag{S10}$$

Notice that $\Sigma_t = \frac{\sigma_{\text{tot}}^2}{\sigma_{\text{env}}^2} > 1$. Next, the phenotypic variance of the population after a reproduction step is given by:

$$\sigma_\theta'^2 = \frac{\sigma_\theta^2\sigma_{\text{env}}^2}{\sigma_{\text{tot}}^2} = \frac{1}{\Sigma_t^2}\sigma_\theta^2 \tag{S11}$$

Therefore, in both equations (S10) and (S11), we see that both the variance and the difference between the mean and optimal phenotypes reduces by the same factor as a result of a reproduction step, that is by $\frac{1}{\Sigma_t}$.

##### Evolutionary dynamics over several generations

Consider a generation as a cycle composed of a fixed number of mutation and HGT steps that occur in a random order, followed by one reproduction step. We wish to calculate the variance phenotype distribution after one generation, given that the initial phenotypic variance of a population is $\sigma_\theta^2(0)$. For this, we first calculate the variance $\sigma_\theta^2$ after $M$ mutation and $H$ HGT steps:

$$\sigma_\theta^2(M+H) = \left((\sigma_\theta^2(0) + m_1 p_m \Delta\theta^2)(1-\frac{p_h}{4}) + m_2 p_m \Delta\theta^2\right)(1-\frac{p_h}{4}) + ..., \tag{S12}$$

where $m_1$ is the number of mutation steps before the first HGT step, $m_2$ is the number of mutation steps between the first and second HGT steps, and so on, such that $m_1 + m_2 + m_3 + ... = M$. After $M$ mutation and $H$ HGT events, there is a reproduction step, after which we can calculate the variance as given by the following:

$$\frac{\sigma_\theta^2(M+H)\sigma_{\text{env}}^2}{\sigma_{\text{env}}^2 + \sigma_\theta^2(M+H)}. \tag{S13}$$

This will be the initial variance of population during the next generation. This gives us a recursion relation, using which we can calculate the steady state phenotypic variance of the population approximately as follows, assuming that $p_m$ and $p_h$ are sufficiently small. The steady state phenotypic variance $\sigma_\theta^2(t\to\infty)$ is given by:

$$\frac{\sigma_\theta^2(t\to\infty)}{\sigma_{\text{env}}^2} = \sqrt{p_m \frac{\Delta\theta^2}{\sigma_{\text{env}}^2} M} - \frac{1}{8}\left(p_h H + 4 p_m \frac{\Delta\theta^2}{\sigma_{\text{env}}^2} M\right) + O(p_h^2, p_m^2 \Delta\theta^4) \tag{S14}$$

This equation shows that the steady state phenotypic variance depends on its HGT rate $p_h$.

##### Determining the population that optimizes fitness

Next, we can calculate the average fitness of the population after $G$ generations. Simulations show that the variances quickly gets to a steady state value, therefore we will assume that it does not change. In a static environment the optimal phenotype remains constant at $\theta_{\text{env}}$. The average fitness of an individual in the population is then given by the following expression:

$$\langle f \rangle = \frac{1}{G}\sum_{i=1}^{G} f_i = \frac{1}{G\sqrt{2\pi}\sigma_{\text{env}}\Sigma_t}\sum_{i=1}^{G} e^{-\frac{(\Lambda_0-\theta_{\text{env}})^2}{2\Sigma_t^{2+4i}}} \tag{S15}$$

where $\Lambda_0 - \theta_{\text{env}} = \frac{\bar{\theta}_p - \theta_{\text{env}}}{\sigma_{\text{env}}}$ is the normalized difference between the mean and optimal phenotypes, $\Sigma_t^2$ is the normalized sum of variances, $\Sigma_t^2 = \frac{\sigma_{\text{env}}^2 + \sigma_\theta^2}{\sigma_{\text{env}}^2}$, and $f_i$ is the average fitness of the population at generation $i$. We assume that after $G$ generations, the environment $\theta_{\text{env}}$ changes. Therefore, a larger $G$ represents a more static environment, while a smaller $G$ a more fluctuating one.

Note that equation (S15) is composed as a product of two terms: (1) $\frac{1}{G\sqrt{2\pi}\sigma_{\text{env}}\Sigma_t}$, which decreases with the phenotypic variance $\sigma_\theta^2$ and is related to the fitness of the population at steady state; and (2) $\sum_{i=1}^{G} e^{-\frac{(\Lambda_0-\theta_{\text{env}})^2}{2\Sigma_t^{2+4i}}}$, which increases with $\sigma_\theta^2$ and is related to the speed with which the population's phenotype distribution aligns with the environment. The competition between these two terms determines the optimum variance in a complicated way: if the variance is reduced, the steady state fitness increases, but it takes more time to reach it. If the variance is increased, the population adapts faster but reaches a lower steady state fitness. This demonstrates the trade-off between speed and fitness that we elucidate in the main text.

In order to determine the optimum variance, it is useful to consider two limits of equation (S15): one when the variance is large (i.e., $\sigma_\theta^2 \to \infty$), and the other when it is small (i.e., $\sigma_\theta^2 \to 0$). When $\sigma_\theta^2 \to \infty$, $\Sigma_t^2 \to \infty$, and therefore in term (2), all the exponents vanish, i.e., $\frac{(\Lambda_0-\theta_{\text{env}})^2}{2\Sigma_t^{2+4g}} \to 0$. Notice that this limit is equivalent to assume that the phenotype distribution is already "aligned" with the environment and therefore has reached its maximum fitness. The average fitness is given by the following expression:

$$\langle f \rangle \approx \frac{1}{G\sqrt{2\pi}\sigma_{\text{env}}\Sigma_t}\sum_{i=1}^{G} 1 = \frac{1}{\sqrt{2\pi}\sigma_{\text{env}}\Sigma_t} \tag{S16}$$

Therefore, when the variance is large, the population's fitness decreases with its variance and is independent of $G$. Because in this limit, the population is already aligned with the environment, the adaptation speed does not play any role and decreasing the variance always increases the fitness as long as this assumption and limit are still valid. The second limit is that for a small variance, e.g., $\sigma_\theta^2 \to 0$ which implies that $\Sigma_t^2 \to 1$, and we can show that the average fitness is then approximately given by:

$$\langle f \rangle \approx \frac{1}{\sqrt{2\pi}\sigma_{\text{env}}} e^{-\frac{(\Lambda_0-\theta_{\text{env}})^2}{2}} + \frac{e^{-\frac{(\Lambda_0-\theta_{\text{env}})^2}{2}}((\Lambda_0-\theta_{\text{env}})^2 G - 1)}{2\sqrt{2\pi}\sigma_{\text{env}}} \frac{\sigma_\theta^2}{\sigma_{\text{env}}^2} \tag{S17}$$

This implies that, if $(\Lambda_0 - \theta_{\text{env}})^2 G > 1$, then, when the variance is small, the fitness increases with the variance. Under this constraint the optimum variance will be $\sigma_\theta > 0$. Otherwise, the optimum variance is $\sigma_\theta = 0$, which corresponds to the highest possible $p_h$.

#### APPENDIX 2
#### MIGRATION FROM AN EXTERNAL POPULATION

While migration between populations is a vast topic that entails many model choices, here we consider a specific scenario, which represents an extreme condition where migration would always be beneficial. Consider two populations: a focal internal population, whose evolution we study in our model, and a second external population, which is a perfectly adapted version of this population. By perfectly adapted, we mean that individuals in the external population are always aligned with the same environment as the internal population, with the same variance as the internal population at steady state. Assume that both populations are otherwise identical, that is they share the same mutation and HGT rates: $p_m$ and $p_h$ respectively. (This is why the steady state variances are the same.) Both populations experience the same environment $\theta_{\text{env}}$. We will assume that individuals always migrate from the external, adapted population to the internal population, a fraction $p_{\text{mig}}$ at every migration step. This is done to maximize the potential benefits of migration from an external population, and maximize the chance that HGT with immigrant donors increases the phenotypic variance of the population. This occurs in two steps, that we will analyze one by one: first immigrants arrive from the external population and change its composition, and second they engage in HGT with members of the internal population. We will calculate the mean and variance of the internal population after these two steps.

First, a fraction $p_{\text{mig}}$ of individuals from the external population migrate to the internal population. The distribution of the internal population after a migration step is given by:

$$(1-p_{\text{mig}})\frac{1}{\sqrt{2\pi}\sigma_\theta} e^{-\frac{(\theta-\bar{\theta}_p)^2}{2\sigma_\theta^2}} + p_{\text{mig}}\frac{1}{\sqrt{2\pi}\sigma_\theta} e^{-\frac{(\theta-\theta_{\text{env}})^2}{2\sigma_\theta^2}} \tag{S18}$$

Using this equation, we can see that after migration, the new mean phenotype of the population is $(1-p_{\text{mig}})\bar{\theta}_p + p_{\text{mig}}\theta_{\text{env}}$. This will affect the fitness of the population by changing the normalized phenotypic difference term in equation (S15) to the environment, as follows:

$$\Lambda_0 - \theta_{\text{env}} = \frac{\bar{\theta}_p - \theta_{\text{env}}}{\sigma_{\text{env}}} \to (1-p_{\text{mig}})\frac{\bar{\theta}_p - \theta_{\text{env}}}{\sigma_{\text{env}}}. \tag{S19}$$

Second, with probability $p_h$ pairs of individuals engage in HGT within this population, causing the changed distribution to follow the expression given by:

$$\mathcal{P}(\theta \text{ after HGT}) = \frac{p_h}{2}\int \frac{1}{2\pi\sigma_\theta^2} e^{-\frac{(\theta_1-\bar{\theta}_p)^2}{2\sigma_\theta^2}} e^{-\frac{(\theta_2-\theta_{\rm env})^2}{2\sigma_\theta^2}} \delta(\frac{\theta_1+\theta_2}{2}-\theta) d\theta_1 d\theta_2 \tag{S20}$$

$$= \frac{p_h}{2}\int \frac{1}{2\pi\sigma_\theta^2} e^{-\frac{(\theta_1-\bar{\theta}_p)^2}{2\sigma_\theta^2}} e^{-\frac{(-2\theta+\theta_1+\theta_{\rm env})^2}{2\sigma_\theta^2}} d\theta_1 \tag{S21}$$

$$= \frac{p_h}{2}\frac{1}{\sqrt{\pi}\sigma_\theta} e^{-\frac{(\theta-\frac{\theta_{\rm env}+\bar{\theta}_p}{2})^2}{\sigma_\theta^2}} \tag{S22}$$

The final distribution after a full HGT step is given by the following expression:

$$(1-\frac{p_h}{2})\frac{1}{\sqrt{2\pi}\sigma_\theta} e^{-\frac{(\theta-\bar{\theta}_p)^2}{2\sigma_\theta^2}} + \frac{p_h}{2}\frac{1}{\sqrt{\pi}\sigma_\theta} e^{-\frac{(\theta-\frac{\theta_{\rm env}+\bar{\theta}_p}{2})^2}{\sigma_\theta^2}} \tag{S23}$$

Therefore, the phenotypic variance after both steps is given by:

$${\sigma'_\theta}^2 = (1-\frac{p_h}{2})\sigma_\theta^2 + \frac{p_h}{4}\sigma_\theta^2 = \sigma_\theta^2(1-\frac{p_h}{4}), \tag{S24}$$

while the mean is given by:

$$\left(1-\frac{p_h}{2}\right)\bar{\theta}_p + p_h\theta_{\rm env}. \tag{S25}$$

We can use these expressions to calculate the optimal HGT rate as a function of the environmental fluctuation frequency at different migration frequencies $p_{\rm mig}$ (Fig. S2). We found that the optimal HGT rate decreased with the environmental fluctuation frequency even when $p_{\rm mig} \neq 0$, but that the magnitude of the decrease weakened with increasing $p_{\rm mig}$. Briefly, this is because even though migration can bring beneficial phenotypes into a population, their phenotype still needs to spread via HGT before the environment changes again. This requires several repeated HGT events from the same, small set of donors, i.e., the newly migrated individuals. This is unlikely in our model, because we assume that HGT occurs between randomly chosen individuals, and in a random direction (an individual can be a donor or acceptor with equal probability).

#### APPENDIX 3
#### COMPARISON WITH DATA

To compare our results with data from natural microbial populations, we used Multi Locus Sequence Typing (MLST) data, which is a common method to estimate recombination rates in bacterial genomes. We used measurements made using the method ClonalFrame developed by Vos and Didelot, with specifically estimates the ratio of the rate of homologous recombination (a common form of HGT) to the rate of mutation in a genome [1]. Briefly, it does this by comparing profiles of variations and substitutions in multiple bacterial genomes which all belong to the same species. This ratio may be interpreted as the relative rate at which HGT occurs in a genome compared with mutation. Note that though it systematically ignores other forms of HGT, and may therefore be an underestimate, we may still use it for comparative studies. For our comparison, we used measurements from $n = 16$ genomes from Table 1 of Vos and Didelot's study, classified by "ecology", or the common habitat in which the genome is found. For static environments, we used all the terrestrial and endosymbiont genomes from the dataset ($n = 8$ genomes; average HGT rate $27.9 \pm 3.2$; error represent standard error of the mean), while for fluctuating environments, we used all the marine or aquatic genomes ($n = 8$ genomes; average HGT rate $1.5 \pm 0.6$).

#### APPENDIX 4
#### COMPARISON WITH OTHER MODELS

We wish to highlight that our unconventional but simple notion of a vector representation of individual phenotypes can be mapped to conventional, but more complex, genome-based population genetic models. Some versions of such models consider the genome as a string of loci and for simplicity, two alleles (say 0 and 1). One can think of the distance between the genomes of two individuals as being analogous to the difference between their phenotypes (or angles $\theta$) in our model. Consider an example with two loci: genotypes 00 and 11 will have the maximum distance, which in our model would correspond to diametrically opposite vectors (say at angle 0 and $\pi$ (180°)). The two other genotypes, 01 and 10, would be equidistant from both 00 and 11, and in our model, can be thought to correspond to two vectors on either side of 0, at angles $\pi/2$ (90°) and $3\pi/2$ (270°) respectively. In the limit of infinite loci, common in population genetics, the distances between genotypes would become continuous, mimicking the idea that the phenotypes in our model take continuous values. In this sense, our vector representation condenses and simplifies the exact details of which genes are different between individuals into an overall distance between their phenotypes.

Additionally, our model is similar in some aspects with the well-known Fisher's Geometric Model (FGM). Both models use the idea of phenotypes (or traits) represented as continuous values, and individuals that possess these traits, as well as the notion that the environment is the optimal phenotype, where the fitness of the individual is maximum. However, FGM studies multi-dimensional phenotypes (more than one phenotype value per organism), while our model, for simplicity, studies one-dimensional phenotypes. Further, unlike our model, trait values in FGM are not periodic, but extend along a large and potentially infinite trait axis. This creates differences in which evolutionary paths populations and individuals can take in the process of adaptation (as discussed above). Finally, in FGM, individuals in a population do not interact, and only mutate independently, whereas in our model, they interact with each other via HGT.

Additionally, despite its simplicity, our vector-based model surprisingly captures several well-known features of evolution that are present in traditional population genetics models. For instance, it exhibits mutation-selection balance, which typifies the steady state of a population without horizontal transfer (Fig. S4). Without horizontal transfer ($p_h = 0$, where our model is still well-defined) a population settles into a steady state with a characteristic variance. This variance depends on the balance between mutation generating less fit individuals and selection removing them from the next generation. Another phenomenon our model captures is Muller's ratchet, which states that less fit mutants inevitably arise in finite-sized populations, even when all individuals start at peak fitness. Our model naturally displays this phenomenon, as the average fitness of individuals whose phenotypes initially align perfectly with the environment gradually decays over time (Fig. S5). Interestingly, in our model, the presence of HGT can partially alleviate this effect (Fig. S5). Such a "rescue" of Muller's ratchet is also observed in other studies which use more complex population genetics models.

Curiously, our model also suggests a link between the recombination-to-mutation ratio and the phenotypic variance of a population. Specifically, on average, the higher the recombination-to-mutation ratio, the lower the phenotypic variance. Since the former is a widely measured ratio for organisms whose genomes have been sequenced, our model also makes another testable prediction, namely that populations with higher recombination-to-mutation ratios should have tighter phenotypic distributions. However, testing this prediction requires deep sequencing and thorough characterization of natural microbial populations, and is outside the scope of this manuscript.

#### APPENDIX 5
#### PARAMETERS USED

| Symbol | Interpretation | Default parameter value |
|---|---|---|
| $N$ | Population size | 5000 |
| $\theta$ | Phenotype of an individual | $0 - 2\pi$ |
| $\Delta\theta_m$ | Magnitude of change in phenotype after a mutation | 0.05 |
| $p_m$ | Probability of mutation (mutation rate) | 0.01 |
| $p_h$ | Probability of HGT (HGT rate) | $0.01 - 0.10$ |
| $R$ | Number of time steps per generation | 1000 |
| $M$ | Number of mutational steps per generation | 100 |
| $H$ | Number of HGT steps per generation | 900 |

### APPENDIX 6
#### SUPPLEMENTARY FIGURES

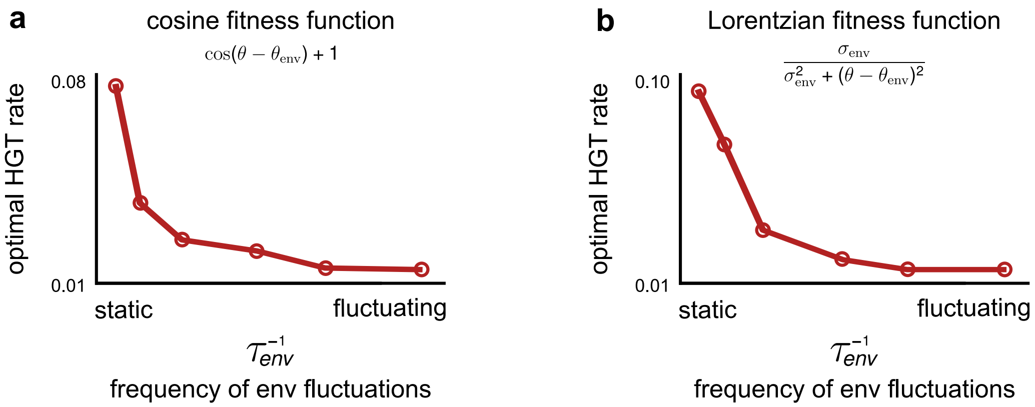


FIG. S1. **Using different fitness functions, as long as they smoothly decrease away from the optimum phenotype, does not affect our main result.** The optimal HGT rate as a function of the environmental fluctuation frequency, $\tau_{\mathrm{env}}^{-1}$; these plots were simulated and derived the same way as in the main text, except instead of using a Gaussian fitness function, we used **(a)** a cosine fitness function (fitness $\propto cos(\theta - \theta_{\mathrm{env}})$) and **(b)** a Lorentzian fitness function (fitness $\propto \frac{\sigma_{\mathrm{env}}}{\sigma_{\mathrm{env}}^2 + (\theta - \theta_{\mathrm{env}})^2)}$). Both fitness functions have one optimum phenotype at $\theta_{\mathrm{env}}$, and the fitness decreases away from it. The width (variance) of the fitness function is given by $\sigma_{\mathrm{env}}^2$.

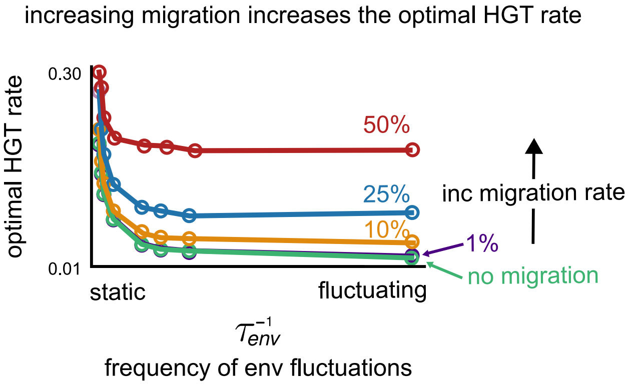


FIG. S2. **Adding migration from an external, well-adapted population does not affect our main result.** The optimal HGT rate as a function of the environmental fluctuation frequency, $\tau_{\mathrm{env}}^{-1}$; these plots were simulated and derived the same way as in the main text, except with an added migration step during a generation. During migration, with a probability $p_{\mathrm{mig}}$ (from 0 to 0.5, marked in different colors), individuals from an otherwise identical external population which is well-adapted immigrate into the internal focal population (aligned with the same environment).

.

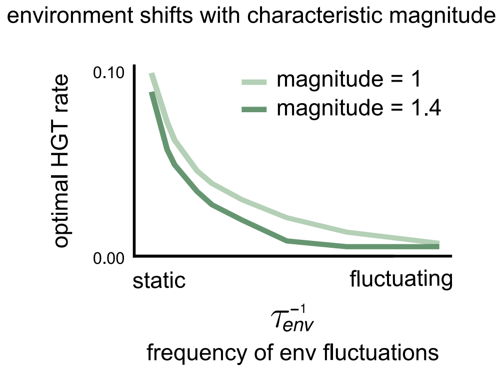


FIG. S3. **Introducing a characteristic magnitude of environmental changes does not affect our main result.** The optimal HGT rate as a function of the environmental fluctuation frequency, $\tau_{\text{env}}^{-1}$; these plots were simulated and derived the same way as in the main text, except during environmental fluctuations, the environment changed by a characteristic magnitude (either by 1, shown in light green, or 1.4 shown in dark green); in the main text, upon fluctuation, the environment changed to a random arbitrary value between 0 and $2\pi$.

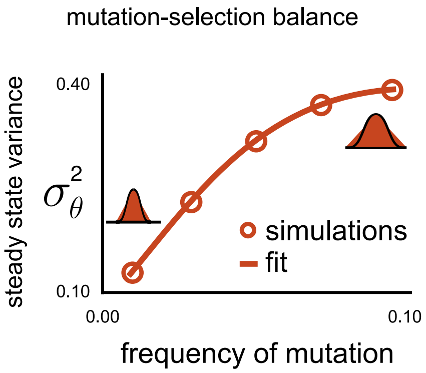


FIG. S4. **Our vector-based model exhibits mutation-selection balance.** The steady state phenotypic variance of a population as a function of its mutation rate ($p_m$ in the model; see Appendix 1) in the absence of HGT ($p_h = 0$). The open circles show results from simulations, while the solid line shows a fit. Populations have a well-defined phenotypic variance, determined by the balance between mutations continually increasing the variance and selection from the environment decreasing it.

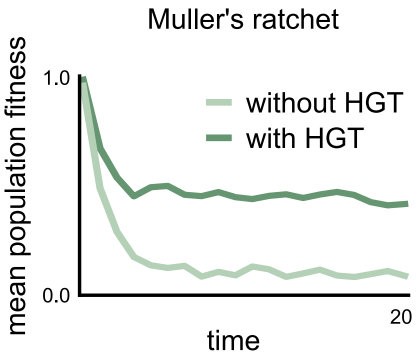


FIG. S5. **Our vector based model exhibits Muller's ratchet.** The average fitness of a population as a function of time (measured in the number of generations), for populations without HGT ($p_h = 0$; light green) and with HGT ($p_h = 0.01$; dark green). Both populations were started with all individuals at the optimal phenotype, thus all being maximally fit. Over time, deleterious mutations accumulate in both populations to lower their average fitness. Having HGT partially weakens this effect, a phenomenon that has been described before as a "rescue" of Muller's ratchet [2].

[1] M. Vos and X. Didelot, The ISME journal **3**, 199 (2009).
[2] N. Takeuchi, K. Kaneko, and E. V. Koonin, G3: Genes, Genomes, Genetics **4**, 325 (2014).
